# LPS-TLR4 pathway exaggerates alcohol-induced acute-on-chronic liver failure via provoking NETs formation

**DOI:** 10.1101/2022.01.25.477506

**Authors:** Yang Liu, Shuo Chen, Jiazhong Wang, Shuo Yu, Xin Zhang, Yiming Li, Gang Cao

## Abstract

Backgrounds: Intrahepatic infiltration of neutrophils is a character of alcoholic acute-on-chronic liver failure (AACLF) and neutrophil extracellular traps (NETs) are an important strategy for neutrophils to fix and kill invading microorganisms. Intestinal bacteria and the gut-liver axis have been thought to play a key role in many liver diseases also including AACLF. However, whether NETs appear in AACLF and play a role in AACLF is still unsure. Methods: WT, NE KO, and TLR4 KO mice were used to build the AACLF model, and the intestinal bacteria were eliminated at the same time and LPS was given. Then the formation of NETs and AACLF related markers were detected. Results: The serum MPO-DNA and LPS concentration was increased in AACLF patients and a correlation was revealed between these two indexes. More intrahepatic NETs formed in AACLF mice by testing MPO-DNA, Cit H3, and NE. These markers decreased with gut detergent and restored markers with gut detergent plus LPS supplement. While NETs formation failed to change with gut microbiome or combine LPS supplement in TLR4 KO mice. As we tested AACLF related characters, liver injury, intrahepatic fat deposition, inflammation, and fibrosis alleviated with depletion of NE. These related marks were also attenuated with gut sterilization by antibiotics and recovered with combined treatment with antibiotics plus LPS. But the liver injury, intrahepatic fat, fibro deposition, and liver inflammation-related markers did show a significant difference in TLR4 KO mice when they received the same treatment. Conclusion: Intestinal-derived LPS promotes NETs formation in AACLF through the TLR4 pathway and further accelerates the AACLF process by NETs.

## Introduction

Alcohol-related injuries are one of the most common causes of preventable illness in the world, leading to 3.3 million people’s death each year and accounting for 6% of all deaths worldwide^1^. Alcohol abuse can insult many target organs and cause multiple system damage. Particularly the liver and gastrointestinal tract are primarily involved in alcohol metabolism, and alcohol abuse causes severe tissue damage. Alcoholic liver disease (ALD) includes a variety of liver diseases such as fatty liver, alcoholic cirrhosis hepatitis (ASH), fibrosis, cirrhosis, and hepatocellular carcinoma. Among them, alcohol-induced acute-on-chronic liver failure (AACLF) or alcoholic hepatitis (AH) is a type of acute on chronic liver disease, the most serious form of ALD, and is characterized by high mortality. According to a report, up to 40% of patients with severe AACLF die within 6 months because there is no effective cure method other than the initiation of prednisolone^2^. Therefore, an intensive understanding of the causes of AACLF is very important for its prevention and treatment.

Intrahepatic infiltration of neutrophils is one of the characteristics of AH^3^. Neutrophils are one of the major effector cells of the innate immune system and have a variety of immune functions such as phagocytosis, degranulation, reactive oxygen species production, while these functions partly rely on or together with neutrophil extracellular traps (NETs) formation and release^4^. Alcohol intake, on the other hand, could cause decreased intestinal motility, overgrowth of intestinal bacteria, and destruction of the intestinal barrier, which can result in the transfer of excess bacteria and their metabolites (like LPS) into the liver^5, 6^. LPS is thought to have the effect of promoting neutrophils to the formation of NET by binding to its receptor TLR4. It has not yet been confirmed whether NET is formed in the liver of AACLF and whether intestinal-derived LPS contributes to the intrahepatic formation of NETs during the condition of AH.

NETs is an extracellular network structure capable of fighting toxic factors and killing bacteria and is mainly composed of extracellular chains of depolymerized DNA and neutrophil granule proteins. These proteins include histones, neutrophil elastase(NE), myeloperoxidase (MPO), cathepsin G, and other more than 30 enzymes and proteins^7^. Although NETs can kill bacteria, fungi, viruses, parasites and further prevent the spread of them and fight against serious infections, the excessive formation of NETs could also lead to exacerbation of inflammation, the development of immune disorders, and damage to adjacent cells^8, 9^. Therefore, NET is considered to a paly role in many diseases such as cardiovascular disease and systemic lupus erythematosus (SLE) ^10^. In the liver, in recent years NETs were also involved in liver ischemia-reperfusion injury, non-alcoholic disease, chemical-induced chronic liver fibrosis, hepatocellular carcinoma et. al^11–13^. However, whether NETs are also related to AACLF is still unsure. Based on the above information, we proposed the following hypothesis that intestinal-derived LPS may promote intrahepatic NET formation via its receptor TLR4 and exacerbate the process of AACLF.

## Method

### Human sample collection

Blood samples were collected from patients with AACLF hospitalized in the Liver disease center at the Second affiliated hospital of Xi’an Jiaotong University. Control samples were collected from patients diagnosed with thyroid nodular goiter and gallbladder stone and without liver disease and alcohol consumption history in the Department of General Surgery, the Second affiliated hospital of Xi’an Jiaotong university before their operation. All the human and animal study was reviewed and approved by the Ethics Committee of Second Affiliated Hospital of the Xi’an Jiaotong University (No. 2018-2115).

### AH Model establishment

The C57 mouse was purchased from the Experimental Animal Center of Xi’an Jiaotong University while NE KO and TLR4 KO mice (C57 background) were purchased from Jackson Laboratory. Mice were kept bred in an SPF animal room with a 12-hour light-dark cycle and not restricted to water and food. Male C57 mice aged 8 to 10 weeks were used for the experiment. During the first 6 weeks, mice were injected intraperitoneally with CCl4 (0.2 ml/kg) dissolved in olive oil (1:4) twice weekly. At week 7, gastric construction catheterization was done by isoflurane-anesthetized according to the method described by Shinji Furuya, and mice were subcutaneously injected with buprenorphine(0.24 mg/Kg) every 12 hours for 72 hours after surgery. At the same time, stop the injection of CCl4 for 1 week. From the 8th week, CCl4 (0.1 ml/kg) was injected intraperitoneally, and meanwhile, a customized animal high-fat diet (Future Biotech, Beijing, China) containing alcohol was injected through the gastric tube. The high-fat diet formula refers to the formula originally described by Thompson. This wine contains corn oil (37% of calories) as fat, protein (23%), carbohydrates (5%), and corn oil as fat (37% of the total). Calories), protein (23%), carbohydrate (5%), etc ^14–17^. Alcohol content in the starts at 16 g / kg / day and increases by 1 g / kg every 2 days until 24 g / kg / day.

### Gut detergent and LPS supplement

Replace mouse drinking water with an aqueous solution containing neomycin (0.3 g / L), ampicillin (0.3 g / L), metronidazole (0.3 g / L), and vancomycin (0.15 g / L) 2 weeks before animal model building. Daily increase antibiotics concentration to reached concentration of neomycin (1g / L), ampicillin (1 g / L), metronidazole (1g / L), and vancomycin (0.5 g / L) and then keep this concentration in drinking water until sample harvest. The stool was collected and tested colony-forming unit (CFU) with blood agar plated to confirm bacteria was removed. In the combined LPS treatment group, LPS (300μg/kg/d) was daily subcutaneous injected.

### Immunofluorescence and immunohistochemistry

Bake the slices in an oven at 60 °C for 30 minutes, then dewax with xylene and sink the slice into the degraded concentration of alcohol. Then place it in a hot antigen recovery solution for 10 minutes. After blocking the antigen with goat serum (DakoCytomation), the primary antibody (1: 100) was added overnight. After washing, the fluorescently labeled secondary antibody (1: 200) was added and washed again, added DAPI. Then, the antifluorescent quencher was dropped and the cover glass was attached. In immunohistochemical staining, the primary antibody was added to the sections after antigen blocking, then the secondary antibody was added and washed, DAB was added for color development, and then the sections were washed and hematoxylin was added. After cleaning, soak in 75%, 85%, 95%, 100% ethanol and soak in xylene. Finally, cover the film for observation.

### Oil red stain

After dehydrating the tissue with 20% and 30% sucrose solutions, the sample is embedded with an OCT embedding and placed in a cryostat to precool. Cut the tissue into 10 μm slices and place it on top of the slices. After soaking in distilled water and 60% isopropanol, soak in oil red dye solution for 5 minutes, in 60% isopropanol, stain with hematoxylin, mount the slide, and observe.

### Determination of serum LPS concentration

LPS concentration was detected using a commercially available color development LPS detection kit (Thermofisher) using the manufacturer’s protocol. Briefly, a 1:50 diluted serum sample (50 μL total) is incubated with 50 μL LAL at 37 ° C for 10 minutes, then 100 μL of chromogenic substrate solution is added 6 minutes later for 37 ° C in the incubation 100 μL. Acetic acid was measured at an absorbance of 410 nm and reported as LPS concentration IU / ml.

### Real-time PCR

Fresh tissue was lysed in Trizol, chromofal was added to separate the layers, centrifugation was performed, the supernatant was added to isopropanol, and centrifugation was performed to collect the precipitate. Then use the DNA-free kit DNAse to remove DANs according to the protocol. Take 1 μg of RNA and apply iScript ™ Reverse Transcription Supermix (Bio rad) to prepare the cDNA. Prepare a 20 μL reaction system using 2 μL of cDNA, primers, iQ ™ SYBR Green Supermix, and water. Comparison of expression target genes mRNA and GAPDH, and relative amounts of gene expression provided for use with the Ct method of ΔΔ. The average expression of the target gene in the control group was set to 1.

### ELISA

Liver samples were weighed and homogenized on ice with cell extraction buffer (Wanleibio. Co., China) containing protease inhibitors. After vortexing the sample for 30 minutes, the sample was centrifuged at 5000 rpm to collect the supernatant and the protein content was quantified by BCA. An assay based on bovine serum albumin. Samples were diluted with PBS to equalize protein concentrations. First, the 96-well plate was coated with a coating solution, and the plate was washed with wash buffer. The wells were then blocked with blocking buffer and standard solution and samples (100 µL) were added to the wells. After incubation, the wells were washed, the detection antibody solution was added, and then the streptavidin-horseradish peroxidase solution was added. Next, the TMB substrate solution was added, and the stop solution was added. Added after 30 minutes of incubation. The absorbance was read at 450 nm. The concentration was determined based on the standard curve.

### Statistics

The results were analyzed by Prism 8.0 software. Numerical data are shown as mean ± standard deviation. The student’s t-test was used to compare the differences between the groups. A P value below 0.05 was considered significantly different.

## Result

### MPO-DNA and LPS are elevated in the serum of patients with acute alcoholic hepatitis

Serum samples were collected from a total of 46 healthy patients and 23 patients with acute alcoholic hepatitis. The levels of LPS and MPO-DNA in the patient’s serum were tested. When neutrophils formed NET, the chromatin DNA associated with other neutrophil proteins was released. So we can determine that the nucleosome is derived from the NETs by measuring the MPO-NA complex ^18^. The characters of clinical data from these patients are shown in Table 1. Due to different disease backgrounds, many differences present between the two groups, like age, gender, ALT, and other test results. Through testing, we found that the serum LPS and MPO-DNA concentrations in acute alcoholic hepatitis patients were significantly higher than the serum concentrations of healthy control people (Fig. 1A, B). More interestingly, we also found that there is a correlation between serum LPS and MPO-DNA concentration in alcoholic hepatitis patients. Given this result of human sample research, we suspect whether NET plays some role in acute alcoholic hepatitis?

**Figure 1.**
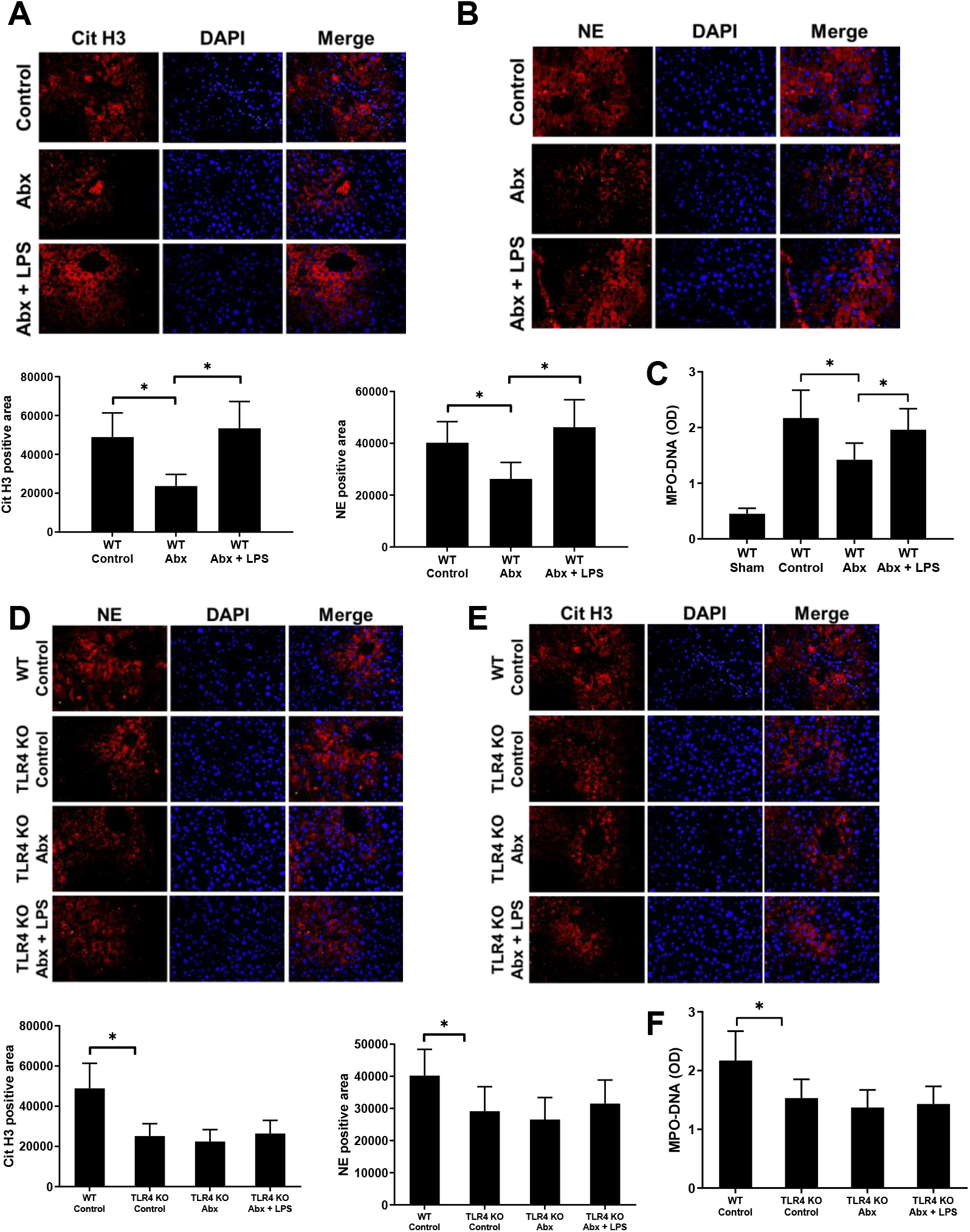
Increased NETs marker and LPS in human AACLF and mice AACLF model. (A) AACLF, patients had an increased serum level of MPO-DNA when compared with control patients without liver disease or alcohol consumption history. (B) Serum LPS concentration was compared between AACLF patients and control patients. (C) Correlation between serum MPO-DNA and LPS concentration in AACLF patients. (D, E)Cit H3 and NE were detected by the immunofluorescence method with the liver sample. (F)Serum LPS concentration from sham and AACLF mice detected by ELISA method. *: *P* <0.05.

### Increased NET formation in mouse AACLF model

Although NETs formed in human AACLF, whether they also formed in rodents is still unsure. So, we built a mice AACLF model as described to figure out the question. This model is considered to imitate a series of human characteristic changes of AACLF such as hepatocyte damage, neutrophil infiltration, intestinal bacteria, and fungi intrahepatic translocation via the portal vein, liver inflammation, and fibrosis^19^. After treating mice with alcohol, the serum ALT concentration and liver TG content were significantly higher than that in the sham group. A large amount of intrahepatic red-stained areas was observed revealed by both Oil-red staining and Sirius red staining. We have also seen many α-SMA and collagen-1a1 positive stained areas in the liver by immunohistochemistry staining. These results suggest that the AACLF mice model was successfully made. Compare with the sham group, the serum LPS concentration was much higher in AACLF mice, and serum MPO-DNA concentration is significate increased in AACLF mice than in sham-treated mice. Intrahepatic fluorescence of NE and CitH3 which was revealed by immunofluorescence staining was significantly higher in AACLF mice than that in sham mice. These results suggest that a large number of NETs formed in the liver after chronic alcoholic liver injury in both humans and rodents. Now we doubt that does NETs relate to AACLF and why NETs formed in this condition.

### Gut microbiota and LPS are important factors for NET formation in the condition of AACLF

Although our previous experiments suggest that NETs are present in the liver of AACLF, what is responsible for this phenomenon is still unsure? To answer this question, a combined model of gut detergent and AACLF was used. The model of gut detergent with a cocktail of antibiotics has been considered to eliminate the gut microbiome effectively and safely^20, 21^. As a result, the intrahepatic fluorescence of Cit H3 and NE was significantly decreased after gut sterilization than control in the AACLF mice model. Serum MPO-DNA concentration was also lower in the antibiotics using group than the no antibiotics control group. This suggests that gut microbiota is a critical factor for NET formation in AACLF. However, this cocktail antibiotics recipe unselectively removed almost all gut microbiome, which substance is the key factor? Lipoteichoic acid, the PAMP of G+ bacteria, is believed to be the reason for the protective effect of intestinal probiotics in various studies. While LPS, the PAMPs of G− bacteria, have been found to have the function which stimulates NETs formation in many different experiments. Therefore, we speculated that LPS probably is the key factor among bacteria and plays a critical role in NET formation in the condition of AACLF. So we give mice with LPS after removal of intestinal bacteria and found that supplement of LPS increased intrahepatic fluorescence of Cit H3 and NE than non LPS supplement mice in the combined gut detergent plus AACLF mice model. Meanwhile, we also tested serum MPO-DNA concentration, and results showed that its level increased in the combined given LPS group than no LPS supplement mice in gut detergent plus AACLF mice.

### TLR4 during alcohol consumption promotes the formation of NET

TLR4 is thought to be the binding receptor for LPS. Many studies suggested that LPS promotes NET formation via TLR4 and further induced tissue damage. Therefore, we speculate whether TLR4 is also involved in liver damage exacerbated by NET during AACLF. A TLR4 KO mice were used to build the AACLF animal model, and results showed that the serum MPO-DNA level of TLR4 KO mice was significantly lower than that of WT mice after alcohol administration. At the same time, the fluorescence of intrahepatic NE and CitH3 in TLR4 KO mice was also significantly lower than the fluorescence of NE and Cit H3 in WT mice. Then we combined the given TLR4 KO mice with antibiotics or antibiotics plus LPS in the model of AACLF and what is interesting is that TLR4 KO mice showed a different pattern compared with WT mice. Our results have shown that there weren’t significant changes of serum MPO-DNA in TLR4 mice either when they received antibiotics or antibiotics plus LPS. Intrahepatic NE and Cit H3 were also detected by immunofluorescent method and we found that, not like WT mice, there wasn’t a significant difference between control, antibiotics, and antibiotics plus LPS group in TLR4 KO mice AACLF model. These results suggest that the intestinal-derived LPS stimulates NETs formation by TLR4 in AACLF.

### NETs is essential for AACLF in mice

Since we found the numerous NETs present in the AACLF both in humans and mice, NETs has been considered to damage adjacent cells in a variety of bacterial and sterile pathological diseases or conditions. Therefore, we speculated that NET may also play some role in the etiology of AACLF. The NE KO mice were used in the study. Neutrophil elastase (NE) could translocate to the nucleus during NETosis and cleaves histones, thereby contributing to chromatin depolymerization^4^. So neutrophils won’t respond to microorganisms and release NETs when NE is absent^22^. In the experiment, we found that NE KO mice had significantly lower serum ALT concentrations than WT mice in the model of AACLF. Liver TG content was also significantly lower in NE KO mice than that in WT mice. By oil-red staining of mouse liver, it was observed that the liver of WT mice had more red-stained areas than NE KO mice. We also tested the mRNAs of lipogenesis-related genes and found that the mRNA expression of ACC1 and FASN in NE KO mice was significantly lower than the expression of WT mice. As for inflammation-related genes, the intrahepatic TNFα, Ccl2, and Cxcl1 mRNA expression levels in NE KO mice were significantly lower than the level of WT mice. We also tested several indicators associated with liver fibrosis. By Sirius-red staining and immunochemistry staining of αSMA and collagen 1a1, it was is shown that WT mice had more intrahepatic positive staining area which indicates more fibro formation than NE KO mice. Meanwhile, the expression of TIMP2 and Collagen1 mRNA which is associated with liver fibrosis was also significantly reduced in NE KO mice than WT mice by qPCR detection.

### Eliminate intestinal bacteria decreased AACLF in mice

As we have demonstrated that NET promotes AACLF, and gut-derived LPS promotes NETs formation, and the gut-liver axis and the intestinal microbiome has believed to be the causes for many alcohols and non-alcohol-induced liver insults. Do we suspect whether gut bacteria and LPS relate to AACLF? We built a combination of intestinal detergent and AACLF model, and results show that antibiotics significantly reduced serum ALT concentration in AACLF. TG levels in the liver were significantly reduced without intestinal bacteria and oil red staining showed significantly reduced intrahepatic red-stained areas in the antibiotics group than that in the control group. The mRNA expression of ACC1 and FASN which is associated with liver tissue lipogenesis was also significate attenuated with the removal of gut microbiota. Expression of liver TNFα, CCL2, Cxcl1 mRNA which are associated with inflammation was lower when mice received antibiotics compared with control. Intrahepatic fibro deposition was also reduced with gut sterilization by detection with Sirius-red staining and staining with αSMA and collagen 1a1. The expression of liver TIMP2 and collagen 1 mRNA decreased with gut detergent compared with the control group.

### LPS is the key for intestinal bacteria insulting the liver in AACLF

Next, we investigated whether LPS is the key to promoting the effectiveness of the intestinal microbiome in AACLF. When LPS was co-administered to antibiotics treated mice, serum ALT concentration and liver TG content were significantly restored compared to antibiotics treated AACLF mice. The intrahepatic fatty deposition was revealed by oil red staining and the result showed that more red staining area was seen after LPS supplement than no LPS supplement in gut sterilized AACLF mice. Intrahepatic fiber which was detected by Sirius-red staining and immunochemistry staining of αSMA and collagen 1a1 was re-increased after giving LPS to gut detergent mice. Liver lipogenesis-related gene ACC1 and FASN, inflammation-related gene TNFα, CCL2, Cxcl1, fibrosis-related genes TIMP2 and Collagen1 also had a higher expression level in LPS plus antibiotics treatment group than only antibiotics treatment group.

### AH Attenuate when mice absent TLR4

Concerning TLR4 playing a key role in intestine LPS induce NETs formation, it is probably also related to the AACLF process. In the study, we found that TLR4 KO mice significantly decreased serum ALT concentration than WT mice. The red-stained areas of the TLR4 KO mice liver were significantly less than WT mice by red oil-red staining, and the liver TG content was also significantly lower in TLR4 KO mice than WT mice. The expression of ACC1, FASN TNFα, CCL2, and Cxcl1 mRNA in the liver of TLR4 KO mice was significantly decreased than in WT mice. Sirius staining and immunohistochemical staining of α-SMA and collagen 1a1 revealed that TLR4 KO mice had less positively stained region than WT mice. At the same time, the liver mRNA expression levels of TIMP2 and collagen1 were significantly alleviated in TLR4 KO mice than those of WT mice.

### TLR4 deficient mice fail to respond to intestinal microbiome and LPS in AACLF

Interestingly, unlike WT mice in the model of AACLF, TLR4 KO mice did not show significant diversity when they removed the intestinal microbiome or supplemented LPS after the gut was sterilized. We tested serum ALT concentration and liver TG content and results showed that there wasn’t a significate difference between the control group, antibiotics group, and antibiotics plus LPS group in TLR4 KO mice of the AACLF model. Intrahepatic red staining areas and ACC1, FASN mRNA expression were the same among these three groups in the TLR4 KO mice AACLF model. Fibrous deposits also didn’t show differences between control, antibiotics, and antibiotics plus LPS group in TLR4 KO mice AACLF model by detecting with Sirius red staining, immunohistochemical staining. Liver fibrosis-related genes TIMP2, Collagen1 together with inflammation-related genes TNFα, CCL2, Cxcl1 mRNA expression kept at the same level when removed gut bacteria and removed gut bacteria plus LPS in TLR4 KO mice.

## Discussion

Human AACLF is main charactered by systemic inflammation, hepatocellular injury, intrahepatic infiltration of neutrophils, and surrounding liver fibrosis. However, most existing animal models of alcoholic liver disease are characterized by minor changes in liver histology, lack of neutrophil infiltration into the liver, and severe pericellular fibrosis ^23^, this limited the study in the field of alcoholic liver disease, especially the AACLF. In this study, we used a model introduced by Shinji Furuya who succeeded in reproducing the characteristics of human AACLF such as more severe liver damage, steatohepatitis, neutrophil infiltration into the liver, liver fibrosis, overgrowth of Escherichia coli and Candida^24^. So, this model is considered an ideal model for AACLF research ^19^.

It has been found that neutrophils increased in peripheral blood and liver tissue tropism of ALD patients and a higher intrahepatic neutrophil level revealed by liver biopsies indicates a poor clinical outcome in AACLF patients^25^. Kupffer cells which are activated after alcohol intake together with damaged liver cells recruit neutrophils to the liver by releasing chemical factors and cytokines. Although their specific mechanism of action is still not fully clear, these infiltrating neutrophils (including immature neutrophils) could lead to liver damage in various ways^26^.

The neutrophil is the main source of ROS. Although ROS formation is an effective sterilization process, excessive or uncontrolled ROS formation can lead to unwanted tissue damage. It had shown that peripheral neutrophils isolated from AACLF or alcohol-related cirrhosis patients had a much higher ROS level compared to neutrophils from healthy donors, which result indicate that more neutrophils were activated in ALD patients^25^. It’s found that serum concentration of some NETs related proteins like lactoferrin, NE, lipoprotein 2 (LCN2) increased in ALD patients which implies more neutrophils may form NETs in this condition^27^. Recently, NET has been also found to induce intrahepatic inflammation in a short-term alcohol intake induced liver injury model^28^, whether NET is present in human and mouse AACLF is still unknown. In the study, we found for the first time that MPO-DNA was elevated in human AACLF through clinical sampling, which suggested the presence of NET in the human body of AACLF patients. However, our result was based on the test of blood sample and could not determine whether NET is located in the liver or some other extrahepatic organs, so we further built an AACLF model in mice and found that a large amount of NETs present in the liver by liver sampling. Then we thought two questions, one is why NETs formed in the liver in this condition, another is that is there some relationship between NETs and AACLF. To figure out these questions, a series of research was conducted.

It’s well known that alcohol could affect multiple end organs (mainly the liver, intestines, and brain). In the gut, it’s found that alcohol could cause an overgrowth of small intestinal bacteria after ingestion. It’s found that ALD patients had lower levels of Bifidobacterium, Lactobacillus spp., Faecalibacterium prausnitzii, Ruminoccoccus spp. as well as Bacteroidaceae, while the proportion of some G− bacteria like Lachnospiraceae was found increased^17, 29^. On the other hand, alcohol could damage the intestinal barrier function and induce intestine hyperpermeability^30^. Although intestinal lumen content of IgA level increased in alcoholics, alcohol consumption impacts several key components of the non-immunologic intestinal barrier and is associated with a decrease of many intestinal tight junction proteins^30^. As a result, excessive intestinal bacteria and its component like LPS migrated into the liver via the portal vein which results were also confirmed in our experiment. In this study, less NET formation was observed with the removal of intestinal bacteria in the process of AACLF. This result suggests that certain microbes or their components may be involved in the formation of NET in this condition. Many in vivo and in vitro studies had proved that LPS, the main component of G- bacterial cell membranes, has been identified as a strong stimulate effect on NETosis^31–34^. While the little study showed that lipoteichoic acid (PAMPs of G+ bacteria) or bacterial DNA had a stimulating effect on NETosis and LTA even shown an inhibitory effect on NETosis in some studies^35^.

When LPS translocates into the liver, it can cause liver damage by various intrahepatic cells through combining with its receptor TLR4. It’s reported that LPS activates Kuffer cells by interacting with CD14 and TLR4 on the surface of the Kuffer cells, then it will release ROS and various inflammatory cytokines and cause damage^36^. LPS could also activate sinusoidal endothelial cells via TLR4 and lead to the release of IL6 and result in liver damage. So, LPS and TLR4 had been considered to have a function to stimulate or regulate NETs formation in many bacterial or non-bacterial related diseases or condition^37–39^. In another double insults model with alcohol and LPS, LPS could also activate hepatic stellate cells and increased deposits of collagen fiber in the liver^36^. However, it is still unclear whether LPS-TLR4 can also insult the liver by stimulation of neutrophils to form NET in the AACLF liver. In the study, we found that LPS restored the NETs formation in gut sterilized AACLF mice, and NETs formation didn’t respond with gut detergent or LPS in TLR4 KO mice. These results suggest that intestinal LPS could stimulate neutrophils to form NETs by TLR4 in AACLF.

Although NET plays a crucial role in innate immunity, excessive NET formation is characterized by promoting inflammation and damaging host cells. Due to these effects, NETs is considered as an underlying basis in many diseases such as systemic lupus erythematosus (SLE), vasculitis, diabetes, thrombosis, lung damage ^40^. In the liver, NET is involved in liver ischemia-reperfusion injury, NASH, hepatocellular carcinoma, etc^11, 12^. More recently, Szabo’s team proved that NET has also been involved in mice liver damage due to short-term acute alcohol intake^28^. In humans, NETs present in AACLF and a higher intrahepatic neutrophil imply a poor outcome of AACLF. In this study, we found that liver damage, intrahepatic inflammation, fat deposition, fibrosis attenuates in the modle of AACLF without NETs which indicates that NETs take part in the process of AACLF. As the gut-liver axis has been considered as the key factor for liver injury in various liver diseases, also including ALD, and we proved that NETs play role in AACLF and gut-derived LPS stimulate NETs formation via its receptor TLR4, so it makes sense that intestinal LPS aggravate AACLF in the study. Although some studies showed alcohol-impaired NET formation in response to antigen^28, 41^, this is probably due to the setting of high circulating lipopolysaccharide (LPS) levels which prime neutrophils and impair their reactivity to further stimuli^13^. However, this study implies that LPS is essential for AACLF present in alcohol consumption at least.

In summary, we can conclude from the study that human and mouse AACLF had an increased formation of NETs in the liver. Intestinal LPS causes the excessive formation of NET in the liver via TLR4, which exacerbates the development of AACLF.

**Figure 2.**
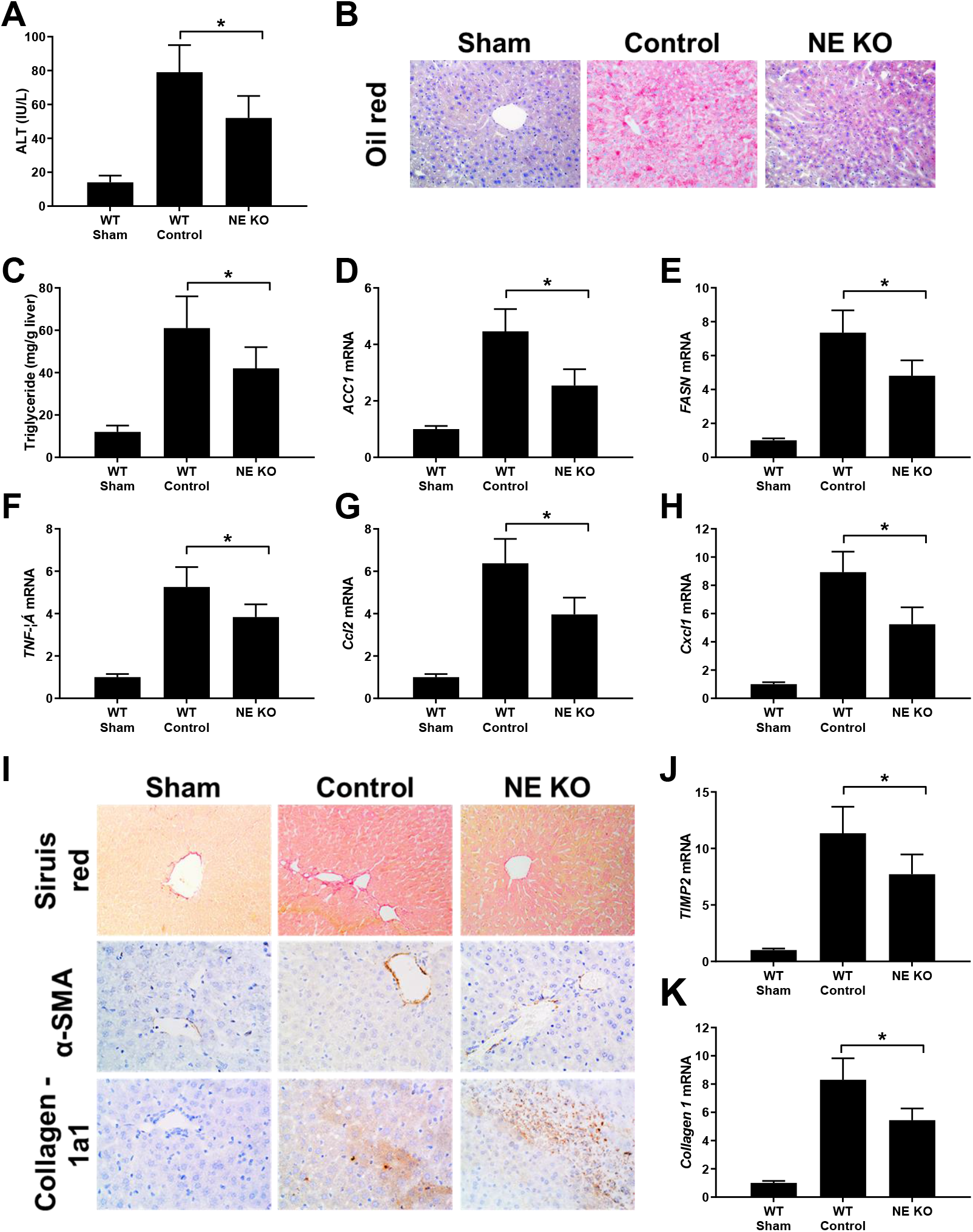
Gut-derived LPS promotes NETosis by TLR4 in the condition of AACLF. AACLF the model was built in WT mice and the mice received gut detergent by cocktail antibiotics or antibiotics plus LPS supplement. Mice were sacrificed after 9 weeks of treatment and liver/serum samples were analyzed by immunofluorescence for Cit H3 and NE (A, B), ELISA for MPO-DNA (C). AACLF, mice model was also built with TLR4 mice and mice received gut detergent by cocktail antibiotics or antibiotics plus LPS supplement. After 9 weeks of treatment, mice were euthanized for sample harvest. Cit H3 and NE were detected by immunofluorescence method with liver sample (D, E) and serum MPO-DNA content was revealed by ELISA method. *: *P* <0.05.

**Figure 3.**
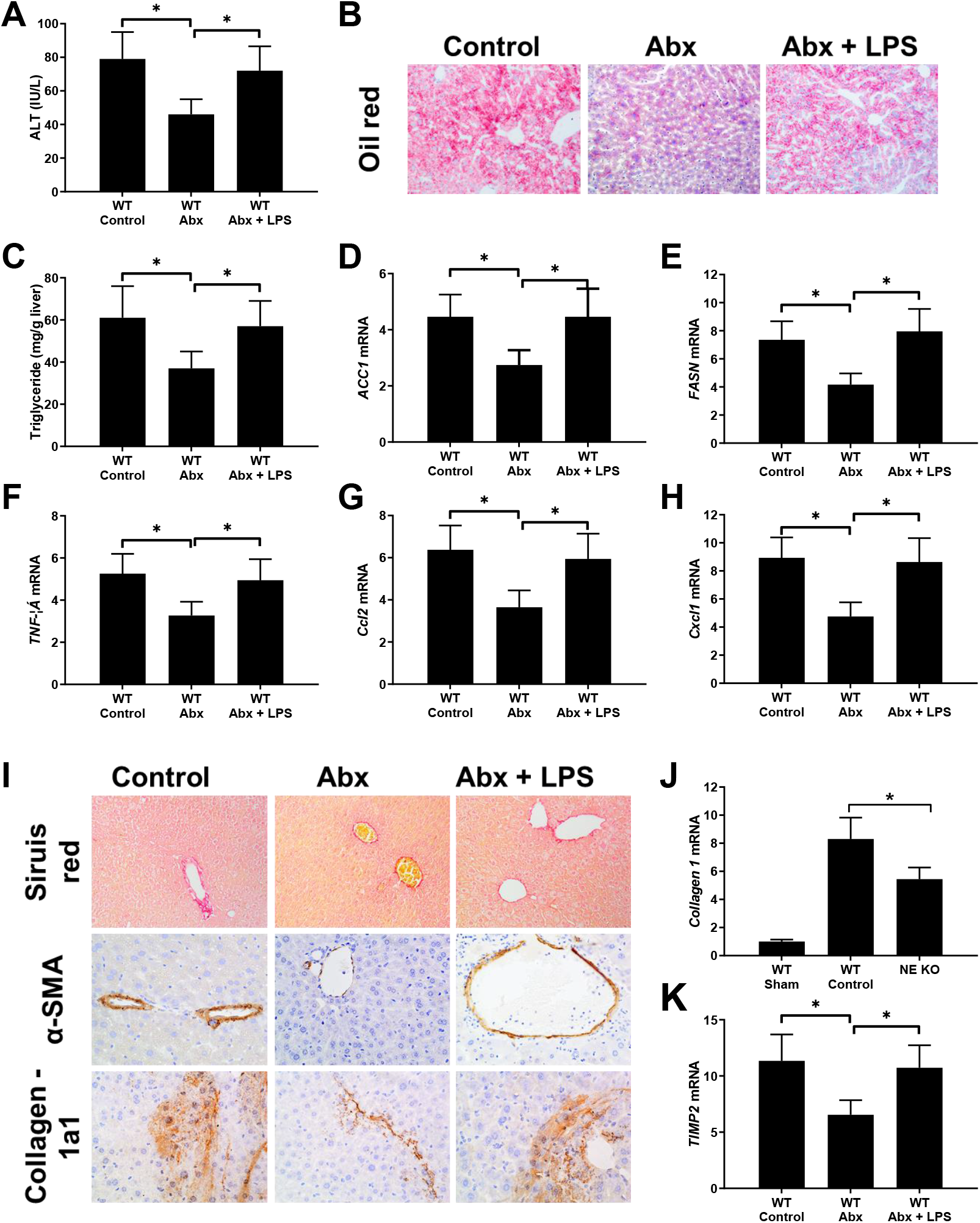
Depletion of NETs decreased liver injury, liver inflammation, fat deposition, and liver fibrosis in mice AACLF model. **AACLF**, the model was built in both WT and NE KO mice. Mice were euthanized at end of model building and liver or serum samples were collected. Serum ALT concentration was revealed by the ELISA method (A). The liver fat deposition was detected by Oil red staining (×20 0) and TG concentration detection (B, C). Liver mRNAs of lipogenesis-related genes (ACC1, FASN) expression was detected by qPCR (D, E). Intrahepatic inflammation was revealed by TNFα, CCL2, Cxcl1 with the qPCR method (F-H). Mice were tested for fibrillar collagen by Sirius red staining (×200) and expression of a −SMA and collagen 1a1 was determined by immunohistochemistry (×400) (I, J). Liver levels of fibrosis-related genes (ACTA-2, TIMP-2, and Collagen-1) were determined by qPCR(J, K).

**Figure 4.**
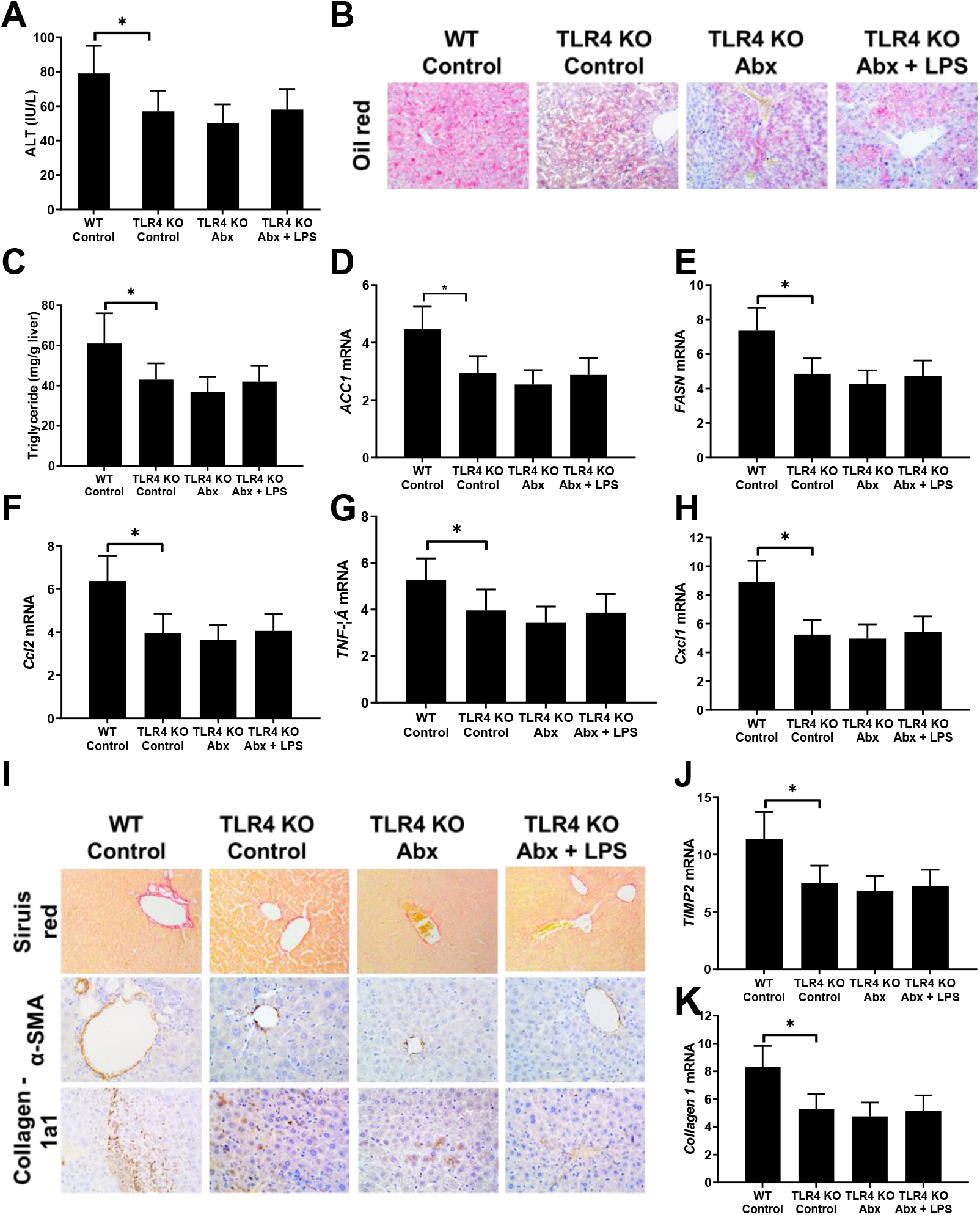
Mice AACLF process decreased without intestinal bacterial and this process recovered after LPS supplement. The intestinal microbiome was sterilized by cocktail antibiotics in the AACLF mice model, and combined given mice with antibiotics plus LPS in another group of mice. Liver and serum samples were detected after sample collection at end of the AACLF model building. Serum ALT concentration was detected by the ELISA method. The intrahepatic fat deposition was revealed by Oil red staining (×200) and liver TG content examination. Liver mRNAs of lipogenesis-related genes (ACC1, FASN) expression was detected by qPCR (D, E). Intrahepatic inflammation was revealed by TNFα, CCL2, Cxcl1 with the qPCR method (F-H). Mice were tested for fibrillar collagen by Sirius red staining (×200) and expression of a-SMA and collagen 1a1 was determined by immunohistochemistry (×400) (I, J). Liver levels of fibrosis-related genes (TIMP-2 and Collagen-1) were determined by qPCR(J, K).

**Figure 5.** TLR4 KO mice didn’t respond with antibiotics and LPS in the AACLF model. AACLF, the model was built in TLR4 KO mice, and mice received different treatment with antibiotics or antibiotics plus LPS. Liver and serum samples were collected at end of the AACLF model building. Serum ALT concentration was detected by the ELISA method. The intrahepatic fat deposition was revealed by Oil red staining (×200) and liver TG content examination. Liver mRNAs of lipogenesis-related genes (ACC1, FASN) expression was detected by qPCR (D, E). Intrahepatic inflammation was revealed by TNFα, CCL2, Cxcl1 with the qPCR method (F-H). Mice were tested for fibrillar collagen by Sirius red staining (×200) and expression of a-SMA and collagen 1a1 was determined by immunohistochemistry (×400) (I, J). Liver levels of fibrosis-related genes (TIMP-2 and Collagen-1) were determined by qPCR (J, K).

